# Activity Budget and Postural Behaviours in Orangutans on Bukit Merah Orang Utan Island for Assessing Captive Great Ape Welfare

**DOI:** 10.1101/2020.12.15.422872

**Authors:** Siti Norsyuhada Kamaluddin, Ikki Matsuda, Badrul Munir Md-Zain

## Abstract

Assessments of the welfare status of captive and semi-captive animals often compare how their expression of natural behaviours differs from that of free-ranging conspecifics. Bukit Merah Orang Utan Island (BMOUI) is the only orangutan rehabilitation and conservation centre in the Malay Peninsula. We recorded and analyzed the activity budget and postural behaviours of orangutans moving freely in the enclosures on BMOUI to evaluate their welfare status. From December 2015 to December 2016, we observed three individuals: an adult male, an adult female and a subadult male, and collected 252 hours of focal data (84 hr/individual). The orangutans’ activity budget was dominated by resting (60%), feeding (13%), playing (14%) and moving (9%). The study individuals heavily relied on the artificial foods (79.2%), and they spent majority of their time on the ground (85.1%) with occasional arboreal observations like using the wooden tree platform or a rope. Despite some significant individual differences, behavioural categories followed a similar trend: resting > feeding > moving > playing, except that the subadult male spent significantly more time playing (35%) than the two adults (3–4%). The most predominant posture was sitting (47.0%), followed by pronograde standing (29.4%), lying (10.5%) and clinging (4.5%). Our results suggest that orangutans on BMOUI engage in less feeding but more resting, and show less postural diversity, than free-ranging individuals. We propose that appropriate interventions to shift activity budgets, especially feeding vs. resting, and postural behaviours of captive orangutans towards those found in free-ranging orangutans might be beneficial for their welfare and survival; however, the conclusions we can draw are limited due to the small sample size, and thus until the captive behaviours of a larger number of orang-utans has been described, these results must be considered preliminary and just a case study.

## Introduction

Conservation programmes have become an integral aspect of zoological management of nonhuman primates, as nearly half of all free-ranging primate species are threatened with extinction due to habitat destruction and poaching (Estrada et al., 2017). The orangutans inhabiting both Borneo and Sumatra are no exception: all are critically endangered. To stop the decline in the number of orangutans it is crucial to establish self-sustaining captive and semi-captive populations, with the ultimate goal of reintroducing individuals to their natural habitats. However, although orangutan rehabilitation is one of the central features in the conservation of this great ape, it is not always successful (Russon et al., 2008).

The ability of orangutans to locate and harvest foods, and food processing skills are considered to be vital for their survival (Descovich et al., 2011). One approach for assessing the psychological aptitudes of captive or semi-captive orangutans is to compare their activity budget with that of free-ranging individuals. It is equally important to postural behaviours, because proficient locomotion skills are critical for safety by allowing efficient resting and foraging high up in trees (Descovich et al., 2011; Grundmann, 2006). However, there is a lack of reports on activity budgets and postural behaviours of captive and semi-captive orangutans based on systematic observations.

BMOUI, located in Perak, Peninsular Malaysia, was established in February 2000 with assistance from the Bukit Merah Laketown Project and the EMKAY Group. In February 2008, the Bukit Merah Orang Utan Island Foundation was incorporated to develop *ex-situ* conservation of orangutans, including rehabilitation and research, and education of the public (Kamaluddin et al., 2018). Here, we report on the activity budget and postural behaviours of three orangutans moving freely in the two enclosures on BMOUI. We include a comparison of the present subjects’ behaviours with those of other captive, semi-captive and free-ranging orangutans, to work towards improving the housing conditions and ultimately the welfare status of the BMOUI orangutans (see Hosey et al., 2009).

## Methods

### Study site and facilities

We conducted observations in BMOUI, in the Malay Peninsula (100° 40.6’ E, 05° 0.50’ N), from December 2015 to December 2016. BMOUI, which comprises an estimated 35 ha. It is covered by secondary tropical forest which consist commercial wood species, e.g., *Dipterocarpus* spp. and *Shorea* spp., and some other cultivated plants (fruits are edible not only for orangutans but also humasn, e.g., *Nephelium lappaceum* and *Bertam* spp.), as the most predominat tree species. Since from the fossil records, it is confirmed that orangutans had historically ranged in the Malay Peninsula (Rijksen & Meijaard, 1999; Tshen, 2016), the natural environment, e.g., vegetation and/or weather, which would partly be comparable to the current distribution habitats of orangutans, exists in the enclosures, suggesting the advantage of the facility of the BMOUI. Four enclosures (A-D), separated by an electric hotwire fence are used as the Orangutan Release Center. The interior of the enclosures contains secondary forest with the addition of ropes, wooden platforms and feeding devices. The orangutans have daily access to the outdoor enclosures from 0900 h to 1700 h, and supplemental feeding is conducted twice per day. Visitors arrive almost daily; they are able to walk through a 100-metre semi-circle steel-fence tunnel that runs through the middle of the enclosures, allowing them to observe the orangutans moving freely in the enclosures (Hayashi et al., 2018).

### Subjects and data collection

There are seven different age–sex classes of orangutans on BMOUI according to Rijksen (1978). The orangutan subjects were placed in two different sized enclosures, i.e., enclosure A (80 m x 180 m): 1 adult male, 1 adult female and 2 subadult females; enclosure B (80 m x 200 m): 3 subadult males. As focal subjects we selected three individuals (Figure 1): an adult male (‘BJ’, 29 years old), an adult female (‘Nicky’, 27 years old, with a 2-year-old infant) in enclosure A, and a subadult male (‘Fatt-Fatt’, 8 years old), in enclosure B. Note that the age information for the adult individuals were estimated upon their arrival to the BMOUI by veterinarians, except the subadult male and the infant of the adult-female subject.

**Figure 1.**
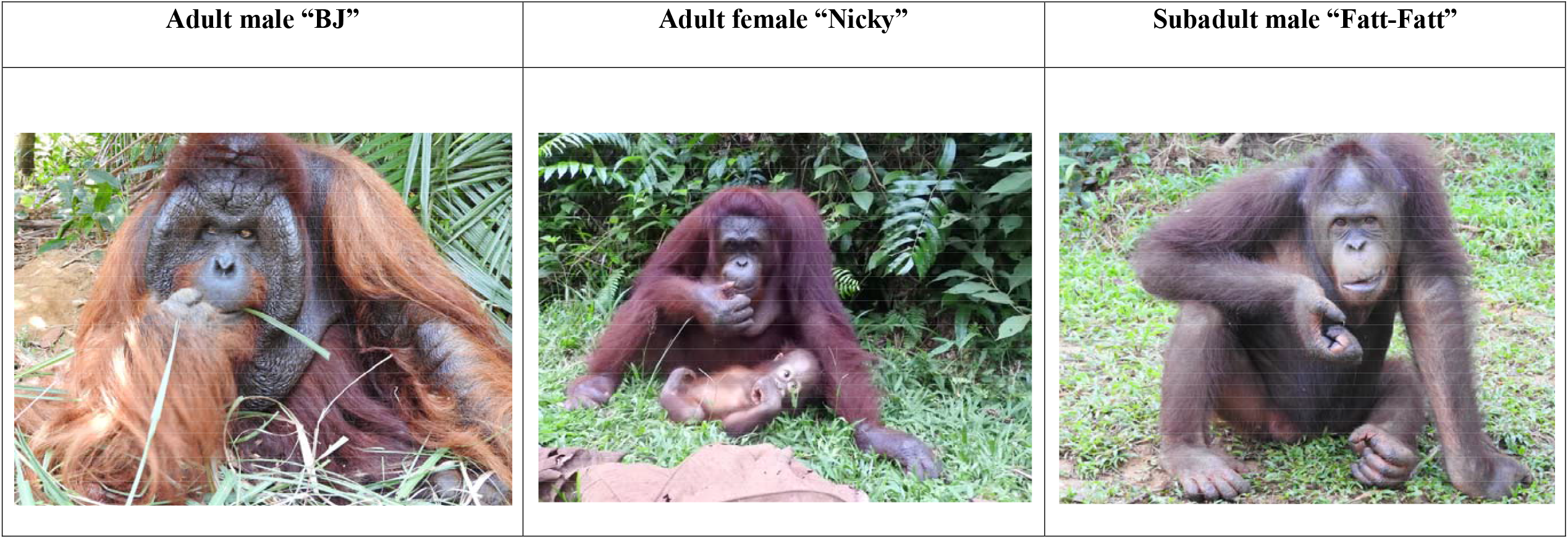
Study orangutans

We followed the three orangutans for 42 days, i.e., 14 days in each individual. Their behavioural data were collected by one of the authors (SNK) continuously during daily observation periods (6 hr: 0930–1330 h and 1430–1630 h), following standardised orangutan data-collection protocols. We used a focal-animal instantaneous sampling technique to record the duration of all behaviours and postures of each individual in 5-min intervals (Morrogh-Bernard et al., 2002).

During focal follows, the observers recorded the time spent in each behavioural activity and the food types eaten (i.e. provided by humans or natural foods). The behaviours were divided into seven categories based on Morrogh-Bernard (2009) and Martin and Bateson (1993): resting, feeding, moving, playing, grooming, and interactions with humans (keepers) including throwing branches, pushing over dead trees, chasing or making kiss-squeaks (Table A1). We also recorded whether the focal individuals were on the ground or arboreal (i.e. on the natural tree/wooden tree platform or using a rope). Postures were divided into eight categories based on Hunt et al. (1996), with some additional definitions added based on Thorpe and Crompton (2006) and preliminary observation by Shariman and Ruppert (2017): sit, lie, cling, pronograde stand, orthograde stand, orthograde forelimb suspend, orthograde quadrumanous suspend and multiposture (Table A2, Figure 2). It should be notable when we analysed the postures, especially while resting and moving in orangutans. Although we defined the stationary behaviour, i.e., resting, as no movement exceeding one minute is counted as a break (Table A1), as a high termination tendency is responsible for many short pauses and a low termination tendency makes for long rests, it was decided to distinguish two categories of stationary behaviour, i.e., “pausing” (shorter than 5 min) and “resting” (longer than 5 min) as followed by the same definition by Sugardjito and van Hooff (1986) and Thorpe and Crompton (2006). For example, when we analysed the siting posture, if the animal engaged in pausing (resting < 5 min), we considered as the siting while moving.

**Figure 2.**
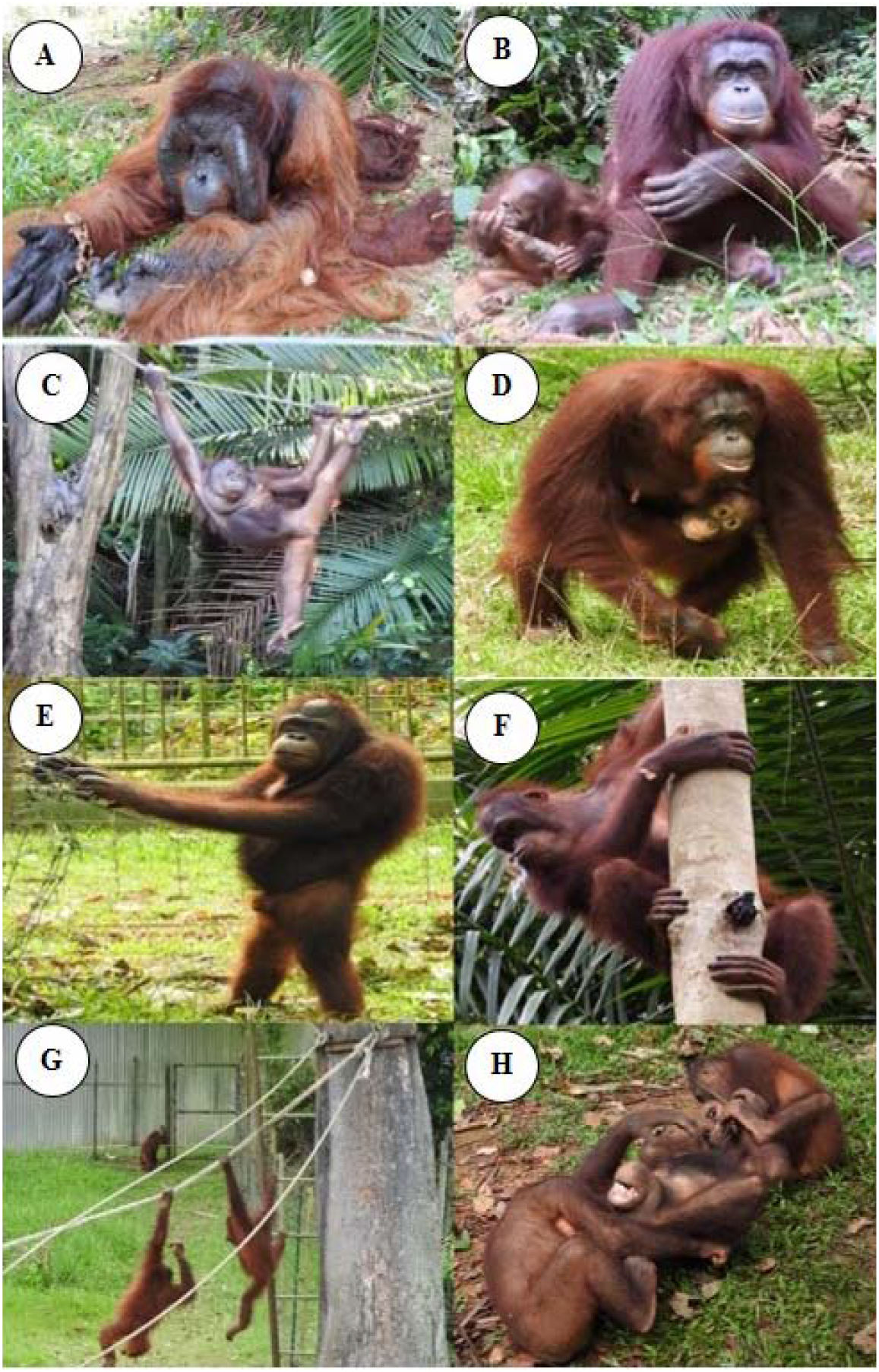
Orangutan postures, a) Lie b) Sit c) Cling d) Ponograde stand e) Orthograde stand f) Orthograde quadrumanous suspend g) Orthograde forelimb suspend h) Multiposture

## Data Analysis

We calculated the activity data with a daily basis, i.e., 14-days data in each individual, and performed Mann–Whitney U tests to compare the mean values (%) of activity budgets, terrestriality, feeding on artificial foods, and postures among the three focal individuals. For each set of hypothesis tests we controlled the overall level of significance at 0.05 using the Bonferroni procedure.

## Results

### Activity budget

We followed three orangutans for a combined total of 252 hours (84 hours per individual). As shown in Table 1 and Figure 3, their activity budgets were dominated by resting (60%), feeding (13%), playing (14%) and moving (9%). The majority of their feeding time was involved in feeding on artificial foods (79.2%). Comparisons of individual activity categories revealed several significant individual differences (Table 2). Time spent resting was highest in the adult male (82%), followed by the adult female (65%), then the subadult male (33%). The adult male spent significantly less time feeding (9%) than the adult female (14%) and subadult male (15%); the adults relied more on the artificial foods (male, mean: 86.7%, SD: 12.2; female, mean: 87.2%, SD: 11.7) compared with the subadult male (mean: 63.6%, SD: 7.80), i.e., adult male vs. adult female: Z= - 0.14 and P = 2.73; adult male vs. subadult male: Z = 3.12 and P < .05; subadult male vs. adult female: Z= -3.03 and P < .05. For all three individuals resting > feeding > moving > playing, except that the subadult male spent significantly more time playing (35%) than the others (3–4%). Noted that subadults mostly fed on leaves, fruits and bark of bertam (*Eugeissona* sp.) and occasionally consumed fruits of cultivated plants and termites as non-artificial foods. Over 70% of the subadult male’s playing time involved social play with other subadults, i.e., playing with other individuals: 71.2%; solitary play: 28.8%. On the other hand, the observed social play with other orangutans was none in the adult male (solitary play: 100 %). Similar tendency was found in the adult female (75.0%), though she engaged in the play with her infant (mean 25.0%, SD: 34.9).

**Table 1.**
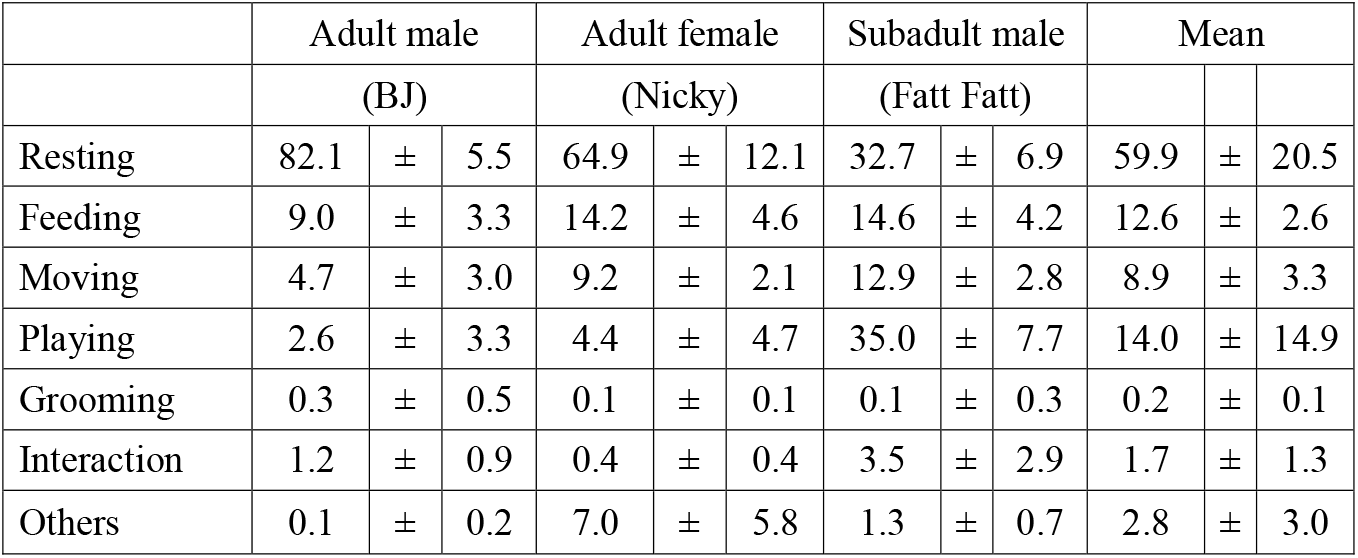
Percentages of activities and for each individual

**Table 2.** Summary of statistical results using a Mann–Whitney U test. A Bonferroni correction was applied for all represented *P*-values. Note that we did not perform the statistical tests for all combination patterns, as it was impossible and/or meaningless for the data including a lot of zeros.

|  | Adult male<br>vs.<br>Female |  | Adult male<br>vs.<br>Subadult male |  | Adult female<br>vs.<br>Subadult male |  |
| --- | --- | --- | --- | --- | --- | --- |
|  | *P* | *Z* | *P* | *Z* | *P* | *Z* |
| **Activity budget** |  |  |  |  |  |  |
| Resting | < .001 | 3.77 | < .001 | 4.50 | < .001 | -4.50 |
| Feeding | < .01 | -2.89 | < .001 | -3.35 | 2.95 | -0.05 |
| Moving | < .001 | -3.49 | < .001 | -4.18 | < .01 | 3.03 |
| Playing | 0.81 | -1.13 | < .001 | -4.51 | < .001 | 4.50 |
| **Posture** |  |  |  |  |  |  |
| **(Resting)** |  |  |  |  |  |  |
| Lying | < .05 | 2.66 | < .001 | 4.27 | < .05 | -2.71 |
| Sit | < .01 | -2.89 | < .01 | -2.99 | 2.62 | -0.18 |
| Cling | 0.06 | -2.32 | 0.08 | -2.20 | 1.13 | 0.92 |
| Pronograde stand | < .001 | -3.45 | < .001 | -4.41 | < .001 | -4.00 |
| Orthograde stand | < .05 | -2.85 | < .001 | -3.46 | 0.31 | 1.66 |
| **(Feeding)** |  |  |  |  |  |  |
| Lying | 2.10 | 0.40 | 1.85 | -0.52 | 0.35 | 1.60 |
| Sit | 2.84 | 0.09 | < .001 | 3.45 | < .001 | -3.45 |
| Pronograde stand | 2.18 | -0.37 | < .001 | -4.04 | < .001 | 3.36 |
| **(Moving)** |  |  |  |  |  |  |
| Lying |  |  |  |  |  |  |
| Sit | 1.44 | -1.44 | 1.44 | -1.44 | 2.19 | -0.15 |
| Pronograde stand | < .001 | 3.74 | < .001 | 4.53 | < .001 | -3.35 |
| Orthograde stand | < .001 | -1.00 | 0.29 | -2.11 | 0.98 | 1.31 |
| Orthograde<br>quadrumanous suspend | < .001 | -3.55 | < .001 | -4.06 | 0.11 | 2.11 |

**Figure 3.**
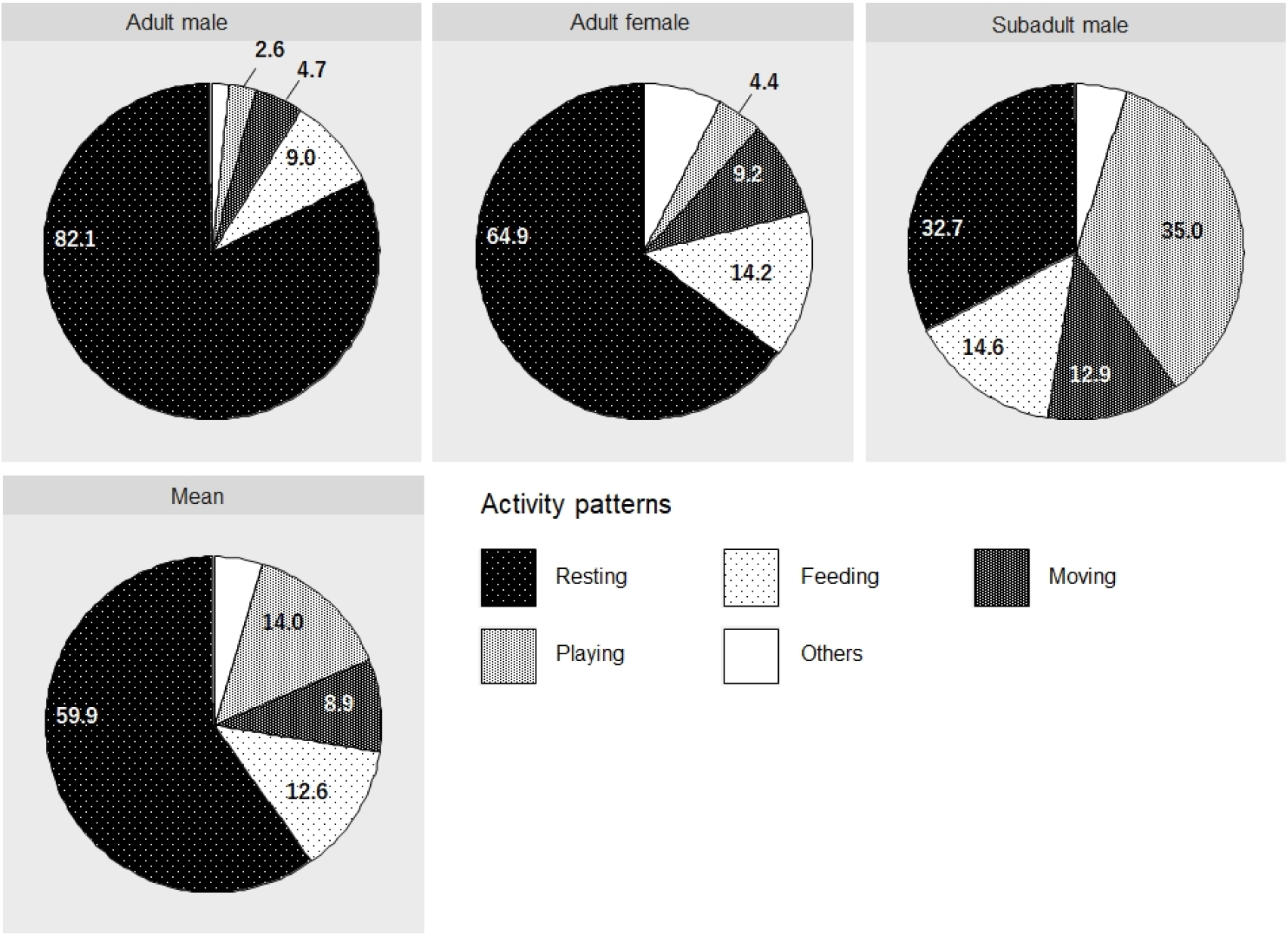
Activity budget for three study orang-utans, i.e., adult male, adult female and subadult male, and their mean

Overall, three orang-utans spent majority of their time on the ground (mean 85.1%±14.7) and were occasionally observed arboreal behaviours using the wooden tree platform or using a rope. The adults (male: mean: 90.0%±16.4; female: mean: 87.8%±15.6) spent more time on the ground than the subadult male (mean: 77.5%±8.52): adult male vs. subadult male: Z= 2.34 and P = 0.06; adult male vs. adult female: Z= 2.15 and P = 0.09; subadult male vs. adult female: Z= - 3.03 and P < .05.

### Postural behaviour

Overall posture (combining feeding, resting and social behaviour) was dominated by sitting (47.0%), pronograde standing (29.4%), lying (10.5%) and clinging (4.5%). Postures while resting, feeding and traveling are shown in Figure 4 (but see Table A3 in details). Sitting was generally the preferred resting and feeding posture; pronograde standing was preferred moving. The adult male sat significantly less (33%) than the adult female (49%) and subadult male (49%) while resting, but he lay more (54%) than the adult female (32%) and subadult male (14%). The pronograde standing posture while resting was most frequent in the subadult male (24%), followed by the adult female (8%) and adult male (4%). Conversely, the pronograde standing posture while traveling was most prevalent in the adult male (98%), followed by the adult female (87%) and subadult male (65%). Orthograde quadrumanous suspension and orthograde forelimb suspension were only recorded during moving (Table 3). Orthograde quadrumanous suspend was significantly less common in the adult male (2%) than in the adult female (10%) and subadult male (13%). Orthograde forelimb suspend was by far the most common posture in the subadult male (21%).

**Table 3.** Percentages of activity budget in comparison to previous studies

| Living-condition | Study site | Activities (%) |  |  |  |  |  |  | *<sup>3</sup>Observation effort | Reference |
| --- | --- | --- | --- | --- | --- | --- | --- | --- | --- | --- |
|  |  | Resting | Feeding | Moving | Playing | Grooming | Interaction | Others |  |  |
| Rehabilitant | Bukit Merah, Perak | 60 | 13 | 9 | 14 | 0 | 2 | 3 | 252 h | This study |
| Rehabilitant | Bukit Merah, Perak | 46 | 20 | 12 | 9 | 8 |  | 5 | 20 h | Hayashi et al. (2018) |
| Rehabilitant | *<sup>1</sup>Bukit Merah, Perak | 52 | 23 | 19 | 3 | 1 |  | 1 | 54 h | Hayashi et al. (2018) |
| Rehabilitant | Bukit Merah, Perak | 62 | 6 | 17 | 12 | 4 |  |  | 30 h | Zaki et al. (2017) |
| Rehabilitant | Bukit Merah, Perak | 33 | 24 | 27 | 9 | 6 |  | 1 | 180 h | C. N. Lee (2003) |
| Rehabilitant | *<sup>2</sup>Meratus Forest, Indonesia | 19 | 58 | 9 | 13 |  |  |  | 740 h | Kuncoro (2004) |
| Rehabilitant | Matang Wildlife Centre, Sarawak | 41 | 23 | 7 | 25 | 2 |  | 2 | 96 h | S. J. Lee (2006) |
| Zoo | Zoo Negara | 58 | 19 | 11 | 9 | 1 |  | 2 | 57 h | May (2004) |
| Zoo | Zoo Taiping | 43 | 21 | 14 | 10 | 12 |  | 1 | 103 h | Wong (2003) |
| Zoo | Singapore Zoo | 50 | 27 | 13 | 8 |  | 2 |  | 192 h | Ting (2011) |
| Zoo | Singapore Zoo | 80 | 9 | 12 |  |  |  |  | 24 h | Matsuda et al. (2017) |
| Zoo | Cleveland Metroparks Zoo, USA | 32 | 15 | 10 | 7 | 3 |  | 3 | 3.5 mo | Cassella et al. (2012) |
| Free-ranging | Batang Serangan, Indonesia | 54 | 24 | 15 |  |  |  | 7 | 2300 h | Campbell-Smith et al. (2011) |
| Free-ranging | Danum Valley Sabah | 34 | 47 | 17 |  |  |  | 2 | 1786 h | Kanamori et al. (2010) |
| Free-ranging | Tanjung Puting, Indonesia | 18 | 62 | 18 | 1 |  |  | 1 | 3805 h | Galdikas (1988) |
| Free-ranging | Mentoko, Indonesia | 45 | 44 | 10 |  |  |  | 1 | 3900 h | Mitani (1989) |
| Free-ranging | Ulu Segama, Sabah | 51 | 32 | 16 |  |  |  |  | 1200 h | Mackinnon (1974) |
| Free-ranging | Suaq Balimbing, Indonesia | 24 | 57 | 18 |  |  |  |  | 11700 h | Fox et al. (2004) |
| Free-ranging | Sungai Wain Forest, Indonesia | 20 | 59 | 14 |  |  |  | 6 | 5 mo | Fredriksson (1995) |
| Free-ranging | Tanjung Puting Indonesia | 16 | 51 | 14 | 2 |  |  | 18 | 349 h | Snaith (1999) |
| Free-ranging | Sabangau, Indonesia | 22 | 61 | 15 | 1 |  |  | 2 | 5502 h | Morrogh-Bernard (2009) |
| Free-ranging | Meratus Forest, Indonesia | 36 | 42 | 17 |  |  |  | 5 | 14 mo | Grundmann (2006) |

\*<sup>1</sup>Behavioral data from semi-captive orangutans using an almost entire island besides BMOUI
\*<sup>2</sup>Behavioral data from rehabilitant orangutans influenced by the rehabilitation program before reintroduced to Meratus Forest, Indonesia
\*<sup>3</sup>Noted that the observation effort in hours would be an important factor influencing the results (activities %), though the number of subjects not in this table may also be an influential factor, especially for the playing category if the subjects include subadults or not.

**Figure 4.**
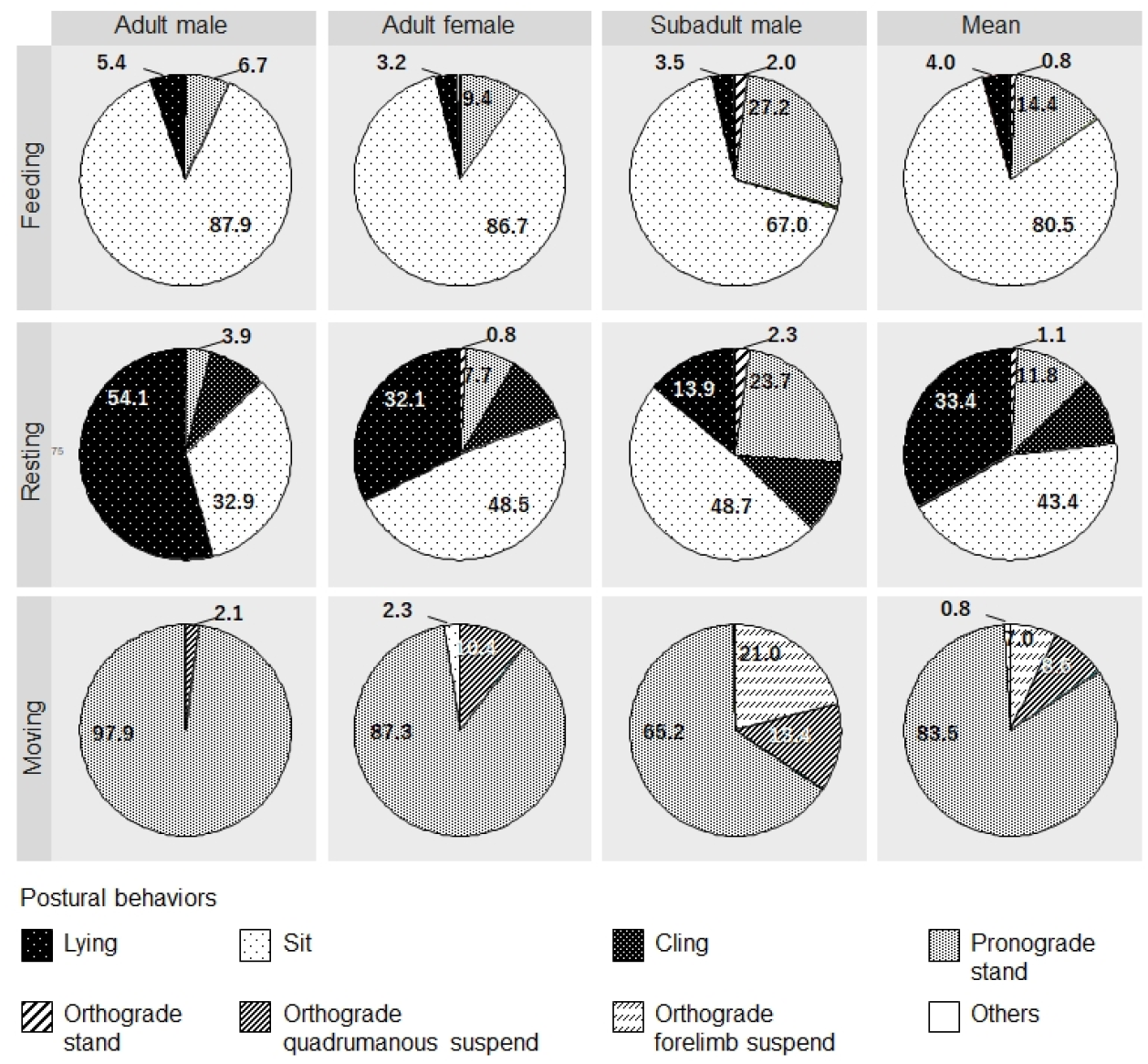
Time allocation patterns of postural behaviour by activity (feeding, resting vs. moving) for each study orang-utan and their mean

## Discussion

### Activity budget

Resting behaviour was one of the major components of orangutan activity budgets in our study, and it was more prominent in the adult male than the others. Since the adult male often monopolised the provided foods at the feeding platform in enclosure A (SNK personal observation), he possibly obtained more foods of higher nutrient content than wild fruits (Nijboer & Dierenfeld, 1996; NRC, 2003; Schwitzer et al., 2009), than the adult female and the subadult male. This might explain why the adult male spent less time feeding and moving (or foraging) and more time spent resting.

Despite being the dominant individual in enclosure B, and therefore able to monopolise provided foods, the subadult male in enclosure B spent less time resting and more time feeding than the adult male in enclosure A. This difference might be related to the greater amount of time spent playing by the subadult, but not by adult even between mother and infant, e.g., < 1% of their waking time in social play (van Noordwijk et al., 2009); indeed, adolescent and subadult males generally engage in more playing behaviours than adults (Galdikas, 1985; Hayashi et al., 2018; Poole, 1987; Schwitzer et al., 2009). In order to spend more time in social behaviours than adults, subadults may need to decrease resting and increase feeding to maintain adequate energy levels. The subadult male also moved around remarkably often, echoing observations of subadult males in the Singapore Zoo (Ting, 2011). Like his zoo counterparts, the subadult male frequently approached tourists, who sometimes gave him food. Zaki et al. (2017) also reported that the amount of time BMOUI subadult males spend moving correlated positively with time spent feeding.

Although resting is the dominant behaviour of both captive and free-ranging orangutans, this may especially apply to captivity (Table 3); free-ranging orangutans generally tend to spend more time foraging, finding, processing and consuming foods, which require both manipulative and cognitive skills (Russon, 1998), and therefore less time resting compared to captive individuals. In fact, the study individuals, especially adults, fed heavily on the artificial foods throughout the study period. In captivity, reduced overall activity levels can lead to weight gain, obesity and undesirable behaviours such as regurgitation and reingestion (Cassella et al., 2012; Pontzer et al., 2010). To reduce the occurrence of such abnormal behaviours and to enhance health and survival, it may be beneficial to intervene to alter the activity budgets of captive orangutans, especially feeding vs. resting, to more closely those of free-ranging orangutans. Providing more diverse food and toy enrichments and devices for cognitive experiments, might play a role in increasing the amount of time foraging and decreasing resting in great apes, e.g., orang-utans (Shariman & Ruppert, 2017) and chimpanzee (Yamanashi & Hayashi, 2011).

Contrary to the free-ranging orangutans which are highly arboreal (Delgado & Van Schaik, 2000), the study captive orang-utans were predominantly terrestrial. Although the recent study reports that terrestrial activity in free-ranging orangutans appears more common than previous anecdotal observations (Ancrenaz et al., 2014), it should not be like our captive orangutans with over 85% of their terrestrial behaviours. Therefore, we believe that this study would be a good example to propose a practical way to alter the behaviors of captive orangutans by introducing enrichment devices such as wooden tree platform and/or ropes, based on the clear contrast between the frequent behavioural patterns exhibited by captive orangutans (more stays on the ground) and ideal one exhibited by free-ranging counterparts (almost completely arboreal). In the meantime, we found the age-related difference in the terrestrial level in this study: adults with larger body size were more terrestrial than the subadult. This may be because larger individual is more dangerous for falling down from high places, and thus the adults would show higher terrestriality, though differing environmental conditions in two different enclosures may simply have created this inconsistency.

### Postural behaviour

Similar to our observations of postural behaviours in this study, other studies have found that most resting time is spent sitting and lying down (Matsuda et al., 2017; Sugardjito & van Hooff, 1986; Thorpe & Crompton, 2006). Sitting while feeding has consistently been more common in other studies (Cant, 1987; Sugardjito & van Hooff, 1986; Thorpe & Crompton, 2006), although pronograde standing was relatively common in our subadult male. This is because he sometimes caught foods thrown by tourists from the ground while in a pronograde standing posture; of course such a situation would not arise in free-ranging conditions.

Pronograde standing was the dominant posture during locomotion in our study, in contrast to previous studies in which sitting was the dominant posture while moving (Thorpe & Crompton, 2006). However, previous studies defined a short rest (< 5 min) as a “pause,” considered as postural mode during traveling; sitting was the dominant posture (Sugardjito & van Hooff, 1986; Thorpe & Crompton, 2006). Nonetheless, as in our study, the second most dominant posture was pronograde stand, suggesting overall similarities in posture, although a direct comparison is impossible.

We rarely observed suspensory postures, especially suspensory quadrupedalism (Niemitz, 2010). These are more common in free-ranging orangutans, not only during locomotion but also during feeding. In fact, suspensory postures accounted for about 24% (Thorpe & Crompton, 2006) and 47% (Cant, 1987) of positional behaviours during feeding. Good locomotion skills are necessary for reintroduced orangutans to stay safe when resting and foraging high up in trees (Descovich et al., 2011; Grundmann, 2006). Hence, modifying captive orangutans’ environments to promote greater diversity in postural patterns and to approach those of free-ranging orangutans, may contribute to their health and survival after reintroduction. One recommendation from this perspective is to suspend food on furnishings including nest boxes, hanging platforms, suspended car tires, trawler nets and wire baskets, and from trees and shrubs to encourage suspensory postures, similar to interventions with other primate species (Britt, 1998).

Lastly, the sort of data presented here was basic, however important to understand the behaviours of captive orang-utans in order to work towards improving their housing conditions and ultimately their welfare. The current information is, nonetheless unfortunately, too limited to differentiate if the differences observed among individuals are not just an outcome of inter-individuals rather than age-sex class differences due to the sample size (only one representative of each age-class). Thus, until the captive behaviours of a larger number of orangutans has been described, these results must be considered preliminary and just a case study. Additionally, in future research, we should look into how a certain kind of enrichment (for e.g., food enrichment) would affect orangutans’ behaviours by examining pre and post enrichment in terms of their activity budget and posture behaviours.

## Authors’ contributions

SNK and BMMZ conceived and designed this study. SNK collected the orangutan behavioural data. SNK and IM analyzed the data. SNK, IM and BMMZ wrote the paper. All authors read and approved the final version of the manuscript.

## Acknowledgements

We thank Bukit Merah Orangutan Island (BMOUI) Foundation, especially Tan Sri Datuk Mustapha Kamal Abu Bakar, Dr. Sabapathy Dharmalingam and BMOUI staff for providing us with the necessary assistance during observations. The authors are deeply indebted to Department of Wildlife and National Parks (PERHILITAN) for providing our research permit. Research methods reported in this research adhered to the legal requirements of Malaysia and were approved by Department of Wildlife and National Parks under research permit (JPHL&TN(IP):100-6/1/14 Jld 2(40). We are grateful to Dr. James Anderson and two anonymous reviewers for their fruitful comments. The authors acknowledge Universiti Kebangsaan Malaysia for providing funding, facilities, and assistance. This research was supported by Grants TD-2014-022, AP-2015-004 and ST 2018-020 (Yayasan Emkay). This study was partly financed by JSPS Core-to-Core Program, Advanced Research Networks (to S. Kohshima) and JSPS KAKENHI (#19KK0191 to IM).

## Supplementary material

**Table A1.** Definition of the activity budget

| Types of activity | Description |
| --- | --- |
| Feeding | Includes all eating activities, such as finding, foraging, manipulating, chewing and swallowing food. Drinking also falls under this category. |
| Resting | Includes all resting activities, such as sleeping, relaxing and watching. The subject does not move much (e.g. observing in a standing, lying or sitting position). |
| Playing | This covers a variety of things, such as playing using mouth and hands. |
| Moving | This includes all types of movements, whether terrestrial or arboreal. Types of body posture include running bipedally or quadrupedally, climbing, brachiating and others. No movement exceeding one minute is counted as a break. |
| Grooming | This activity involves clearing the hair and scratching or dressing any part of the body by hand, foot or mouth. |
| Interaction with human | This activity involves throwing branches, pushing over dead trees and chasing or making kiss-squeaks towards human. |
| Others | Includes all behaviours and acts not listed above, such as dropping to the ground from the rope or tree, urination, nursing a baby and making a nest. |

Source: Morrogh-Bernard et al. (2002) and Martin and Bateson (1993)

**Table A2.** Definition of the postures

| Type of postures | Description |
| --- | --- |
| Lie | Almost all the orangutan's weight was supported on its torso, and it was in a horizontal or reclining position. |
| Sit | Almost all the orangutan's weight was supported on its rear end, and the upper body was upright. |
| Cling | Almost all of the orangutan's weight was supported by hanging with two hands. |
| Pronograde stand | The orangutan's weight was separated equally between its hands and feet while on a horizontal substrate. |
| Orthograde stand | The orangutan's weight was separated equally between its hands and feet. |
| Orthograde quadrumanous suspend | Hanging with two hands and feet on supports angled at $\geq 45^\circ$. |
| Orthograde forelimb suspend | Forearm suspensory posture from tree branch to tree branch. |
| Multiposture | Includes all types of locomotion above, especially when playing alone or with other orangutans. |

Source: Hunt et al. (1996), Thorpe and Crompton (2006) and Shariman and Ruppert (2017)

**Table A3.**
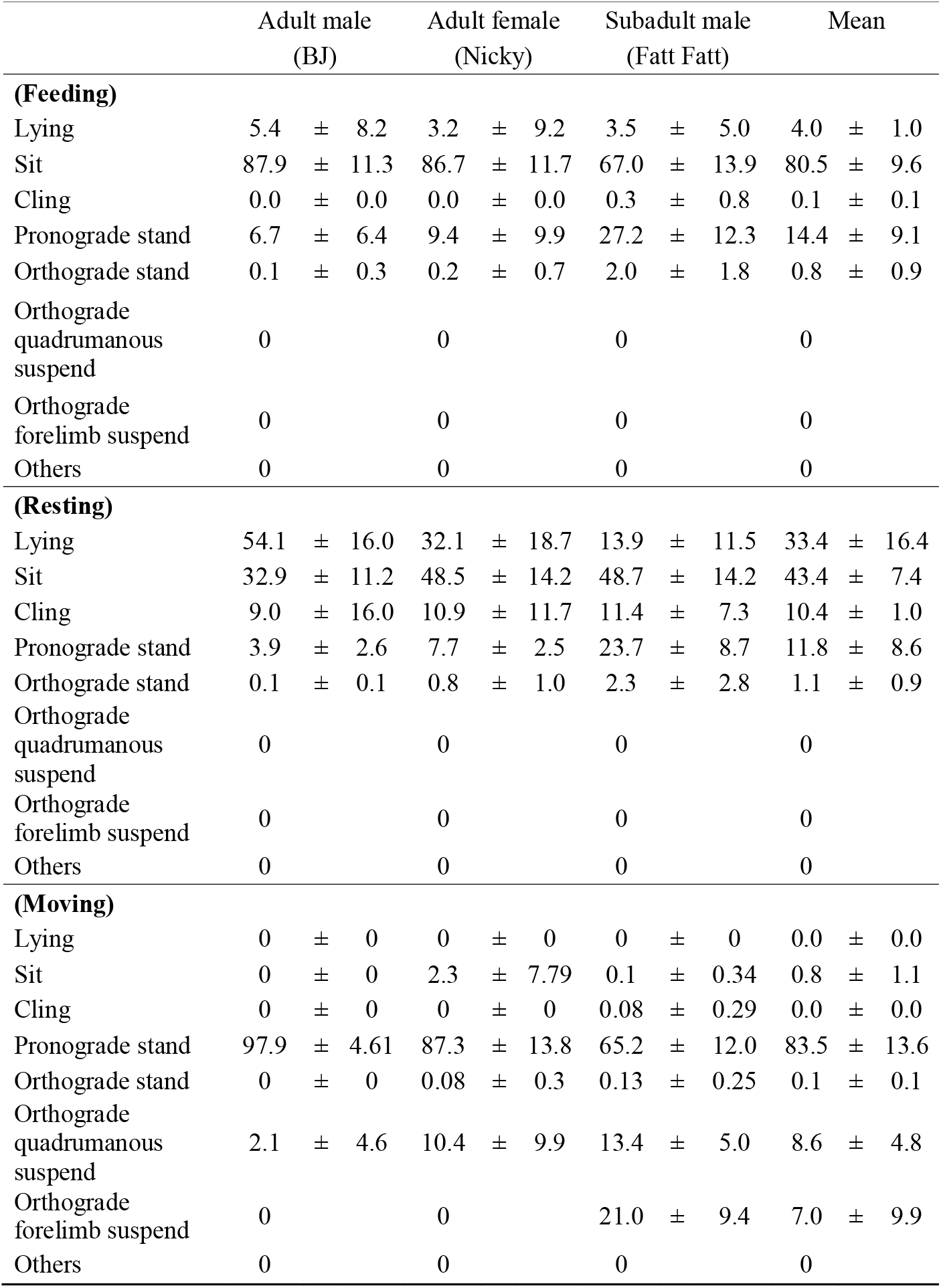
Postural behaviour by activity (feeding, resting vs. moving) for each individual

